# Integrated assessment of structure and dynamics of solid proteins

**DOI:** 10.1101/2022.10.20.513076

**Authors:** Benedikt Söldner, Kristof Grohe, Peter Neidig, Jelena Auch, Sebastian Blach, Alexander Klein, Suresh K. Vasa, Lars V. Schäfer, Rasmus Linser

## Abstract

Understanding macromolecular function, interactions and stability hinges on detailed assessment of conformational ensembles. For solid proteins, accurate elucidation of the spatial aspects of dynamics at physiological temperatures are limited by the qualitative character or low abundance of solid-state NMR internuclear distance information. Here, we demonstrate access to abundant proton-proton internuclear distances for integrated structural biology and chemistry with unprecedented accuracy. Apart from highest-resolution single-state structures, the exact distances enable molecular dynamics (MD) ensemble simulations orchestrated by a dense network of experimental inter-proton distance boundaries gathered in the context of their physical lattices. This direct embedding of experimental ensemble distances into MD will provide access to representative, atomic-level spatial details of conformational dynamics in supramolecular assemblies, crystalline and lipid-embedded proteins, and beyond.

---

Access to site-resolved structural data is usually a precondition for detailed understanding of bio-macromolecular function. Elucidation of key biological and biophysical properties ranging from intermolecular interactions and recognition to enzymatic activity, regulation, and allostery, however, also requires insights into different details of atomic motion. NMR spectroscopy provides access to various aspects of structural dynamics under close-to-physiological conditions. Much in contrast to many other structural-biology techniques providing only ground-state structures, NMR parameters are shaped by additional conformations that are populated at room (or body) temperature and thus can also contribute to the molecular properties via the underlying dynamic conformational ensemble. In solution, molecular dynamics (MD) simulations are regularly augmented by experimental NMR restraints, which are translated into (system-specific) pseudo-potentials that are added to the regular protein force field. These experimentally derived bias potentials can be implemented in the MD simulations via ensemble-averaging approaches that obey the maximum-entropy principle.^[2]^ In addition to chemical shifts, accurate structural restraints (e. g., exact NOEs) are available that permit reconstruction of dynamic ensembles with great detail.^[3]^ Complementing the class of soluble or solubilized proteins and increasingly aided by cryo-EM tomography for information on the nm to µm scale,^[4]^ solid-state NMR (ssNMR) grants access to structural and motional features of proteins in a solid lattice, including supramolecular assemblies, fibrillar or membrane proteins. With the onset of direct proton detection, owing to fast magic-angle spinning (MAS) and/or partial deuteration, ssNMR has seen a revolution with respect to its associated technical possibilities and feasibility for biological studies, with a particular prospect for high-molecular weight targets.^[5]^ Structure elucidation from proton-detected solid-state NMR has initially been hinging on amide and methyl groups, whose contacts are sparse for perdeuterated proteins and have limited structural assessment to a reasonably defined backbone fold.^[1, 6]^ Fully protonated proteins at MAS rates exceeding 100 kHz, by contrast, now open the door to a dense network of internuclear distances. The restraints are an order of magnitude more numerous than for deuterated proteins, but associated, as of now, with severe shortcomings regarding a quantitative interpretation, which compromises the value of established structure elucidation approaches.^[7]^ In the solid state, for detailed elucidation of dynamic ensembles within crystalline proteins, supramolecular assemblies, fibrils, and membrane proteins, a similar impact of concatenation of experiment and simulation via the maximum-entropy principle can be expected as for solution NMR. However, due to the sparseness (e. g., isolated REDOR^[8]^ spin pairs) or rather qualitative character (e. g., current homonuclear recoupling techniques^[1b, 1c, 7a, 7c, 9]^) of distance restraints in modern approaches using fast magic-angle-spinning, prospects have been limited to chemical-shift anisotropy and dipolar-coupling-based implementations in oriented membrane protein samples.^[10]^ Opposing the limited perspectives prevailing for the large range of solid proteins, here we demonstrate integrated assessment of structure and dynamics via experimental data from fast-MAS NMR, leveraged by a dense network of accurate proton-proton distances made attainable.

In solid-state NMR, the buildup of coherence-driven magnetization transfer usually behaves similarly to a linear combination of Bessel functions of the first kind.^[11]^ Here, to revisit the potential of such data for detailed assessment of structure and dynamics, we recorded and assigned 3D time-shared ^15^N/^13^C-edited RFDR magnetization buildups^[1b]^ at 110 kHz MAS on a micro-crystalline, fully protonated (^13^C/^15^N) sample of the chicken *α*-spectrin SH3 domain in a time-dependent fashion (Fig. 1A, see details in SI). In total, 1033 unambiguously assigned peaks were obtained, including 274 cross peaks from bidirectional transfers, 485 cross peaks from unidirectional transfers, and 274 diagonal peaks. For generation of exact solid-state distance restraints, we adapted the eNORA framework^[12]^ to the specific behavior of solid-state NMR magnetization transfers, in particular tailored fitting of magnetization buildup and correction for dipolar truncation and spin diffusion. Bearing advantages over numerical simulations, an analytical function for fitting recoupling curves in a powder sample (i. e., with a distribution of angles β between *B*_*0*_ field and inter-spin vector according to random molecular orientations) was derived by Mueller using a linear combination of Bessel functions of the first kind (*J*_*α*_ with order *α*),^[11]^ whose series expansion can be truncated practically to *k* ≤ 5 elements,

**Fig. 1:**
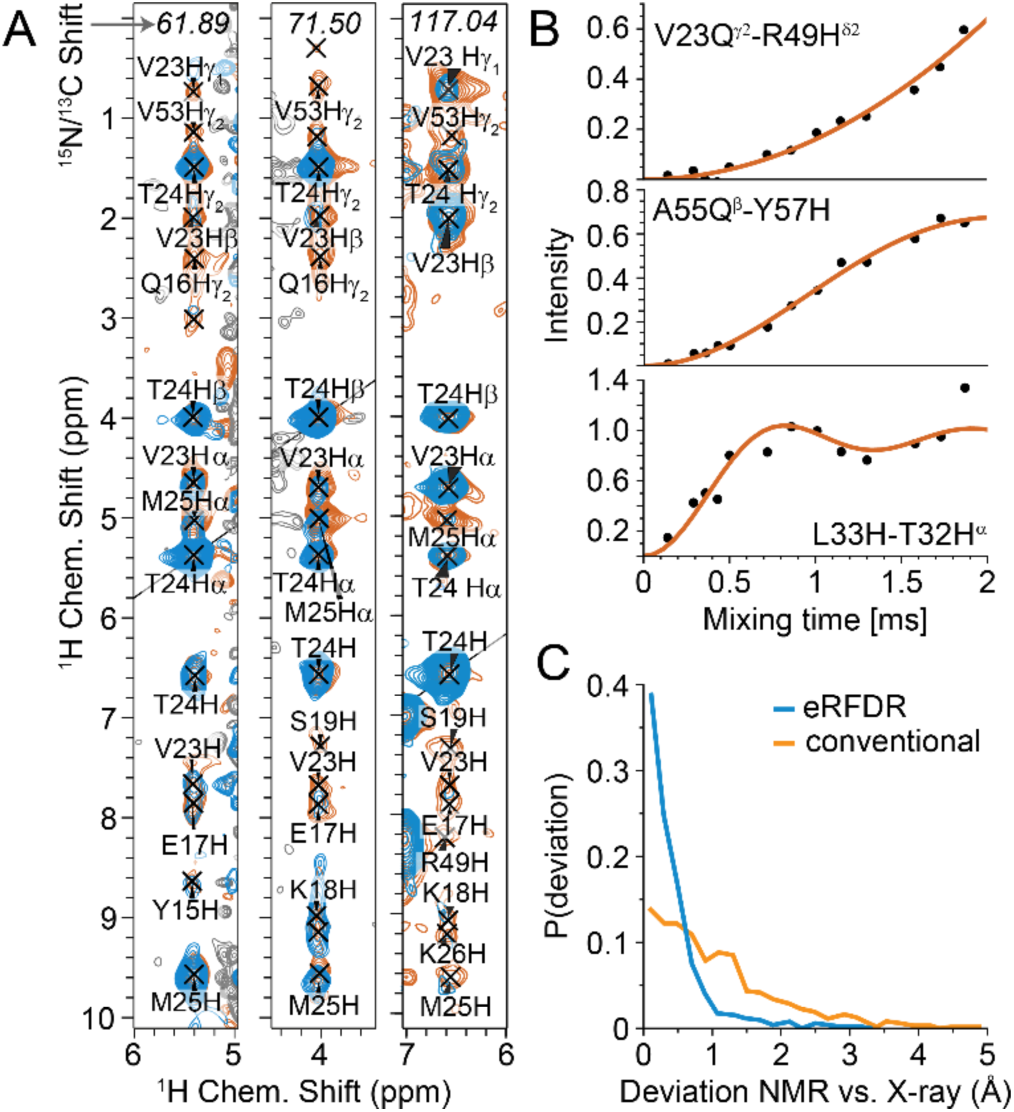
Generation of accurate solid-state NMR distance restraints. **A)** Exemplary strips of a time-shared experiment (diagonal at T24 H^α^, H^β^, and H^N^) for a fully protonated sample of the SH3 domain of chicken *α*-spectrin at 110 kHz MAS (blue: 0.288 ms mixing time, orange: 1.008 ms). Instead of these 3D data, increasing peak dispersion by a fourth dimension is possible.^[1]^ **B)** Buildup curves (slow, intermediate, and fast) and their fits for exemplary transfers (with distances of 6.4, 4.8, and 2.1 Å from top to bottom). **C)** Statistics of the accuracies obtained for distance restraints, shown as deviations to the corresponding distance found by crystallography.

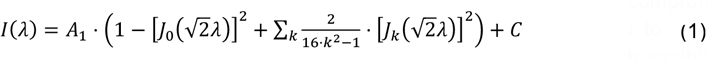

where *I* is the intensity as a function of mixing time, *A* is a vertical scaling factor (amplitude), *λ* is the argument of the Bessel function, and *C* is an optional offset factor (not used within this study). The effective behavior of homonuclear dipolar recoupling and coherence-driven transfers is reasonably similar to the REDOR case (compare Fig. 1B).^[13]^ Hence, the argument *λ* can be constructed as *λ* = *D* · *τ*_*mix*_ from the buildup time *τ*_*mix*_, modulated by the interaction strength *D*, which in an ideal two-spin case is directly proportional to the dipolar-coupling constant. The distance *r* is then obtained via

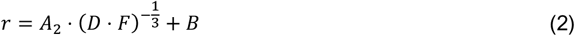

with the correction factor *F* combining spin diffusion and dipolar truncation, and user-defined values for the slope *A*_2_ and the *y*-intercept *B*, both calibrated with respect to a preliminary starting structure or known distances, e. g., between geminal protons. The scaling factor of the apparent interaction strength is not known beforehand but merges into a global fit parameter *A*_2_. Even though the procedures for fitting, extraction of distances from the fitted interaction strengths, as well as the developed methods for correction of spin diffusion and dipolar truncation (see below and the SI for details) are heuristic solutions, the close correspondence of the distances determined from the corrected interaction strengths and the true distances are apparent in Fig. 1C (blue curve).

Direct interpretation of magnetization buildups is compromised by the fact that the magnetization transfer from spin *I* to spin *S* is modulated by nearby third spins (e. g., dipolar truncation) as well as spin diffusion. The latter one is heuristically taken into account by assuming a merely additive effect to the observed magnetization flow, as has been done in the framework of NOE buildup in solution.^[14]^ To obtain correction factors for dipolar truncation, a simulation-based method hinging on nearest-neighbor interactions was developed (see details in the SI). In brief, 17500 theoretical 3-spin scenarios of RFDR transfers from *I* to *S* with a third spin *K* with different geometries, in addition to a simple 2-spin buildup for each *I*-*S* distance, were simulated in SIMPSON^[15]^. All synthetic buildups were fitted in analogy to the experimental data, and for each geometry, a correction factor was derived as the ratio between the apparent interaction strength *D* obtained in the 2-spin simulation and the fit value *D* obtained when fitting the output of the 3-spin simulation. These correction factors, applied based on similarity between simulation geometry and spin environment of the site in a template structure (see below) as well as the strength of distortion comparing different third spins, enabled association of experimental buildups with relevant two-spin distance values.

As correct cross peak assignment can be efficiently obtained from iterative assignment procedures^[16]^, distance restraints were benchmarked here for a series of 15 3D time-shared, ^15^N/^13^C-edited RFDR spectra of different mixing times. With the increased dispersion of 4D spectra^[1]^ the benefits of full protonation could be exploited even further, implying fewer overlaps and hence further improvements of data quality but necessitating more measurement time. As a template structure^[12]^ for correction of spin diffusion and dipolar truncation, we either used (1) the X-ray structure 2nuz (as a prove of principle and for demonstrating the achievable distance restraint accuracies when used as an input for NMR-restrained ensemble MDs), (2) a structure obtained from conventional ssNMR structure calculation (see the SI) based on cross-peak volumes of a single RFDR spectrum (for restraint accuracies obtainable when no X-ray, cryo-EM or AlphaFold^[17]^ structure is available), and (3) an incorrectly folded starting structure (for assessing the error tolerance of the method for correction of spin diffusion and dipolar truncation towards a worst-case scenario, see Table S1). Table S1 also shows that spin diffusion correction is the correction of choice for the deuterated sample, whereas the combination of both corrections yields best results for fully protonated samples. In addition, Table S1 and Fig. S2 compare results for calculation based on only 5 (instead of the excessive 15) RFDR spectra, in which case quantitative restraints are still obtained with only minute decrease of the structural statistics.

For the protonated sample, given the high number of protons available in the side chains, 718 of 759 buildups could be fitted, or 25.6 per residue on average (only including assignable residues, 7 – 62). Among these, 98 peaks were discarded due to unreliable fitting (See the SI for details.), 204 peaks could be included as bidirectional restraints, with magnetization transfer found and quantified both for *i* → *j* and *j* → *i*, and 416 peaks were used as unidirectional restraints (Fig. 2 and Table 1). In contrast, for the deuterated and back-exchanged sample, only 52 peaks could be included as bidirectional restraints and 67 peaks as unidirectional ones, which is only 18 % of the restraints of the protonated case.

**Table 1:**
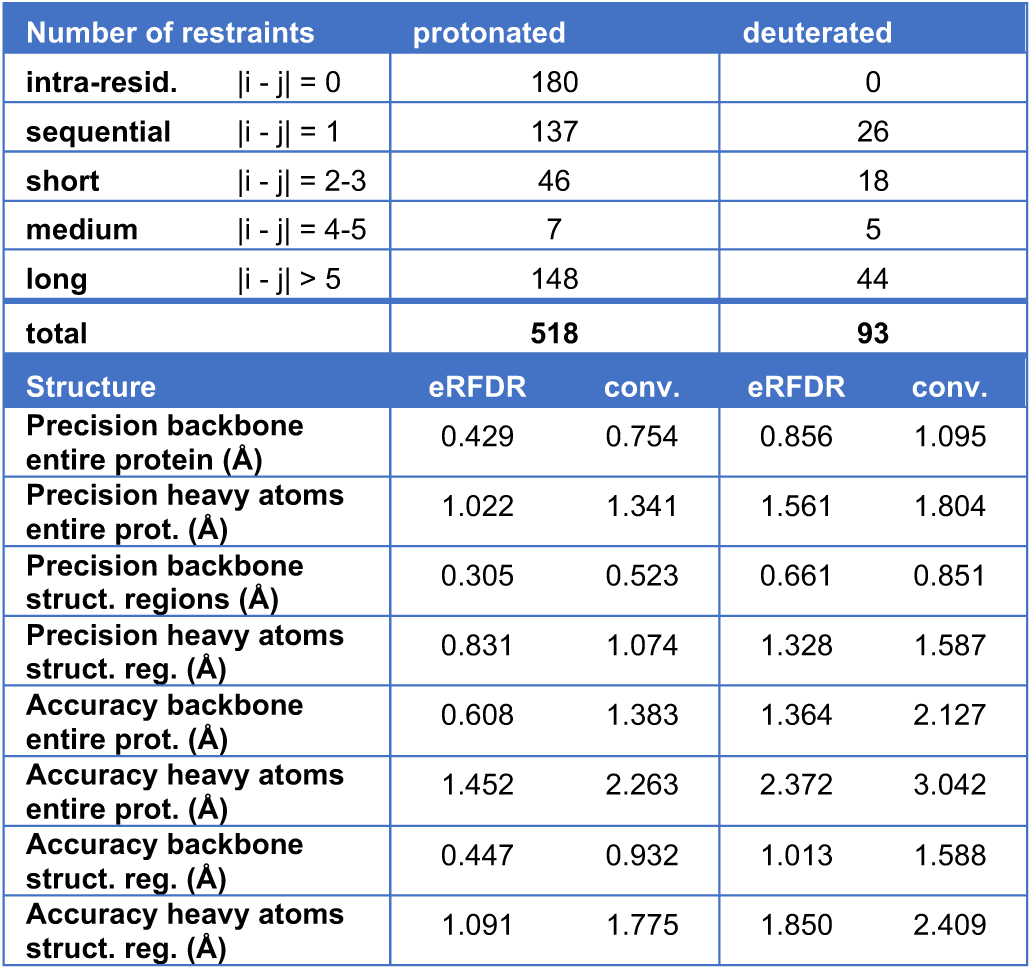
Structural statistics, shown for the entire protein and only for structured regions, i. e., excluding residues 17-23 (RT loop), 36-41 (n-SRC loop), 47-48 (distal loop), and termini, using the combined (spin diffusion and dipolar truncation) correction method with 2nuz as template.

**Fig. 2:**
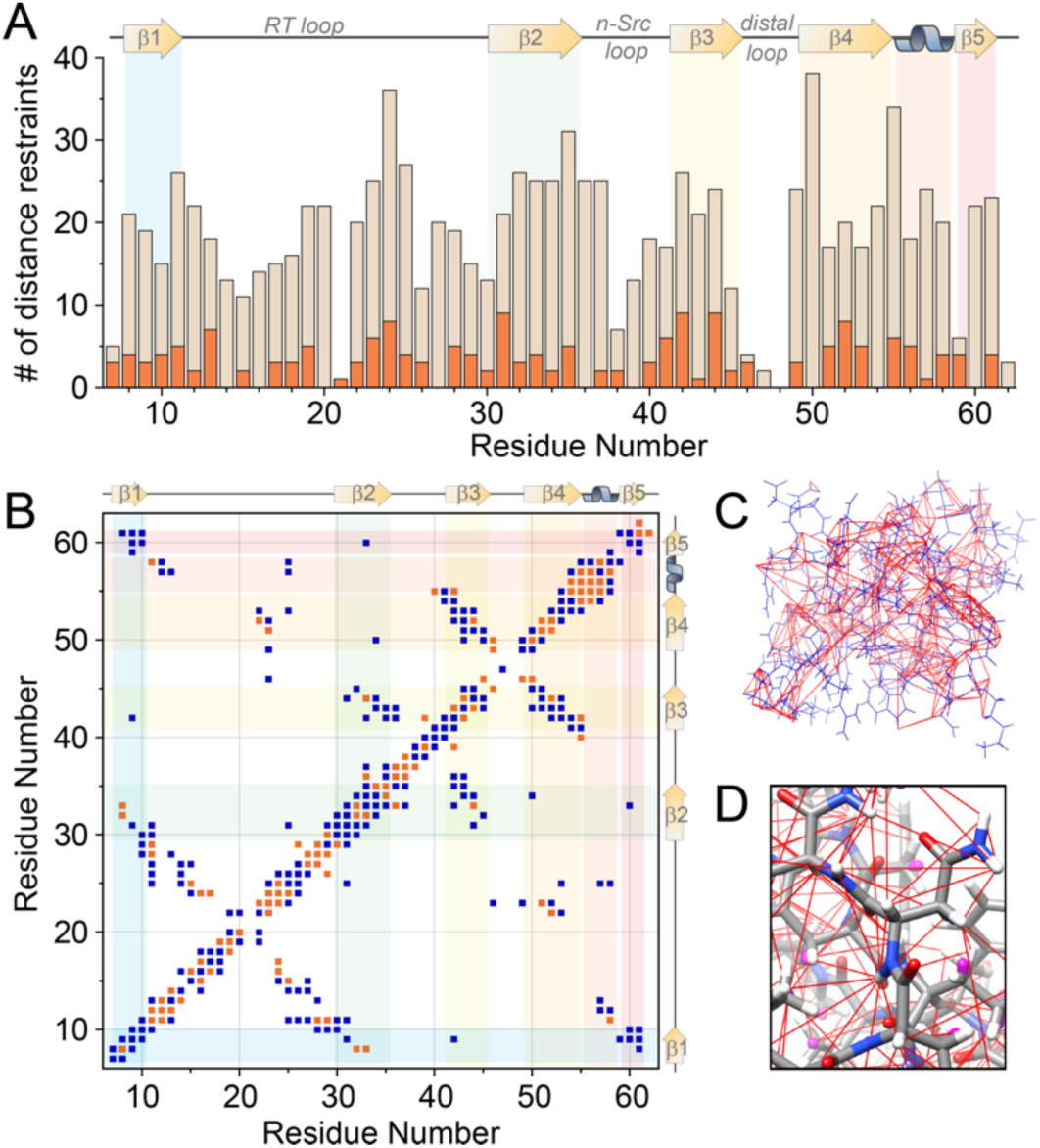
Quantity of the obtained restraints and their connectivities. **A)** Number of distance restraints per residue (ochre: fully protonated sample, orange: deuterated and proton-backexchanged sample). **B)** Contact map with uni-(blue) and bidirectional (orange) distance restraints. (Only one dot is shown per pair of residues even for multiple contacts via different protons.) **C)** and **D)** Depiction of the coverage of obtained distances, forming a dense network of restraints.

At first, in order to assess to which extent the utilization of accurate distance restraints can improve regular structure determination in solid-state NMR using either protonated or deuterated and ^1^H^N^ back-exchanged samples, structure calculations (Fig. 3) were performed with ARIA^[16]^, as laid out in more detail in the SI.

**Fig. 3:**
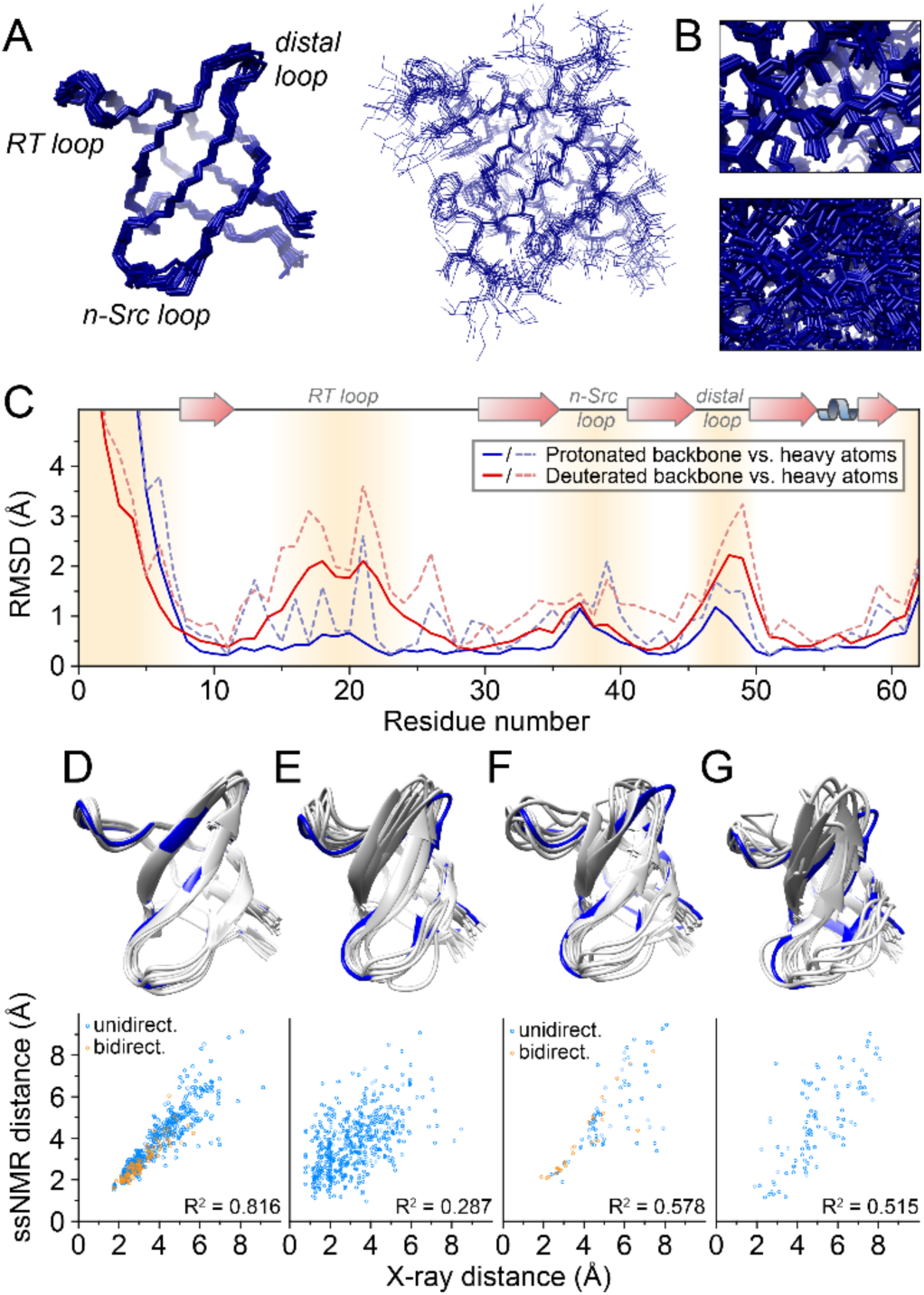
Single-state structure calculation from fully protonated samples via accurate distance restraints. **A)** Backbone structural ensemble and overlay of these structures showing all heavy atoms. **B)** Visual comparison of structural definition of this structure elucidation (zoom-in on the top) with the ensemble resulting from a deuterated/back-exchanged sample, assessed using the same procedure (bottom). **C)** Structural precision as a function of residue for this structure elucidation compared to the calculation derived from a deuterated/back-exchanged sample. **D-G)** Structure calculations from fully protonated (D and E) or deuterated and proton back-exchanged samples (F and G), via eRFDR restraints (D and F) or conventional restraints (E and G), with their corresponding distance correlations to X-ray data (bottom row). eRFDR restraints were determined using the combined (spin diffusion and dipolar truncation) correction method using the X-ray structure PDB 2nuz as template.

Indeed, structure calculation resulting from the high abundance of exact distance restraints yields very high precision with a root mean square deviation (RMSD) of 0.43 Å/1.02 Å (backbone and all-atom, respectively), determined over all residues from 7-61 (or even 0.30 Å /0.83 Å determined over the secondary-structural elements only, see Table 1). The zoom-in in Fig. 3B as well as the residue-wise comparison in Fig. 3C demonstrate the improvement of structural precision with regard to a deuterated/back-exchanged sample under otherwise comparable conditions (backbone and all-atom precision of 0.86 Å and 1.56 Å, respectively). Similarly, comparison of the structural ensemble with the X-ray structure 2NUZ yields a backbone RMSD of 0.45 and 1.01 Å for protonated and deuterated/back-exchanged structure, respectively, for the secondary-structural elements (see Table 1).

More importantly, as even modern state-of the art MD simulations are limited in accuracy for assessing dynamic structural ensembles via MD as a stand-alone technique due to inaccuracies of the force fields used and incomplete conformational sampling, combining MD simulations with experimental NMR data (either by *a-posteriori* reweighting methods or by restrained-ensemble techniques) has become a gold standard of integrated structural biology in solution. The availability of a dense network of exact solid-state NMR distances now enables restrained-ensemble MD simulations for *solid* protein preparations. Such integrated process bears important prospects for detailed assessment of conformational ensembles of the various targets whose overall molecular weight or solubility excludes solution NMR. The maximum-entropy principle is implemented in the MD ensembles by considering only the ensemble average for the comparison with the experimentally derived restraints (which are also ensemble averages). Importantly, the lattice and intermolecular contacts in ssNMR samples are often unknown and challenging to implement in the MD. In the presence of the experimental data, however, restrained-ensemble MD simulations can be carried out for the monomeric protein in solution, whereas the ssNMR restraints now guide the conformational sampling to regions of the energy landscape that are in line with the experimental conditions. The MD ensemble comprised 16 replicas, which were subjected to bias potentials on the proton-proton distances according to the experimental eRFDR restraints from the protonated sample. The simulations were performed for 160 ns with the Amber ff99SB*-ILDN protein force field^[18]^ together with the TIP3P water model^[19]^ (see SI for more details). The number of distance restraint violations is very small (15 out of 518 restraints are violated on average, with an average absolute violation (over all 518 restraints) of < 0.001 nm, sum of all violations 0.388 nm). In Fig. 4, the root-mean-square fluctuations (RMSFs) of the atoms around their mean position from the hybrid NMR/MD assessment are plotted, reflecting atomic-resolution dynamics in the backbone (solid lines) and the entire protein including side chains (dashed lines). In fact, the eRFDR-restrained MD ensemble shows highly similar, but slightly lower sequential RMSFs than the corresponding unbiased MD simulations, with an average decrease of 6.2 and 6.4 % for backbone and all-heavy-atom fluctuations, respectively. The same applies for RMSDs with respect to the X-ray structure (Figs. 4). Despite the tight restraints, the differential behavior of the individual atoms and the relative mobility of different structural elements are highly consistent (correlation coefficient of 0.999 and 0.996 for backbone and all-heavy-atom fluctuations, see also Fig. S8), demonstrating that an artificial “over-restraining” to a static structure is avoided. Conversely, the close agreement found between the experimental and computational results and the very minor restraint violations also validates the MD simulation on its own. It is gratifying to see that the force fields and the amount of conformational sampling applied are appropriate to realistically capture the structural dynamics of the protein. Most interestingly, the data confirms the high similarity of dynamics for this protein domain in a crystal and in solution, complementing previous experimental studies^[3c, 20]^ with atomic-resolution spatial details. By contrast, larger and more flexible proteins are expected to be more sensitively impacted by their lattices, which can be brought to light by the integrative modelling approach outlined in this work.

**Fig. 4:**
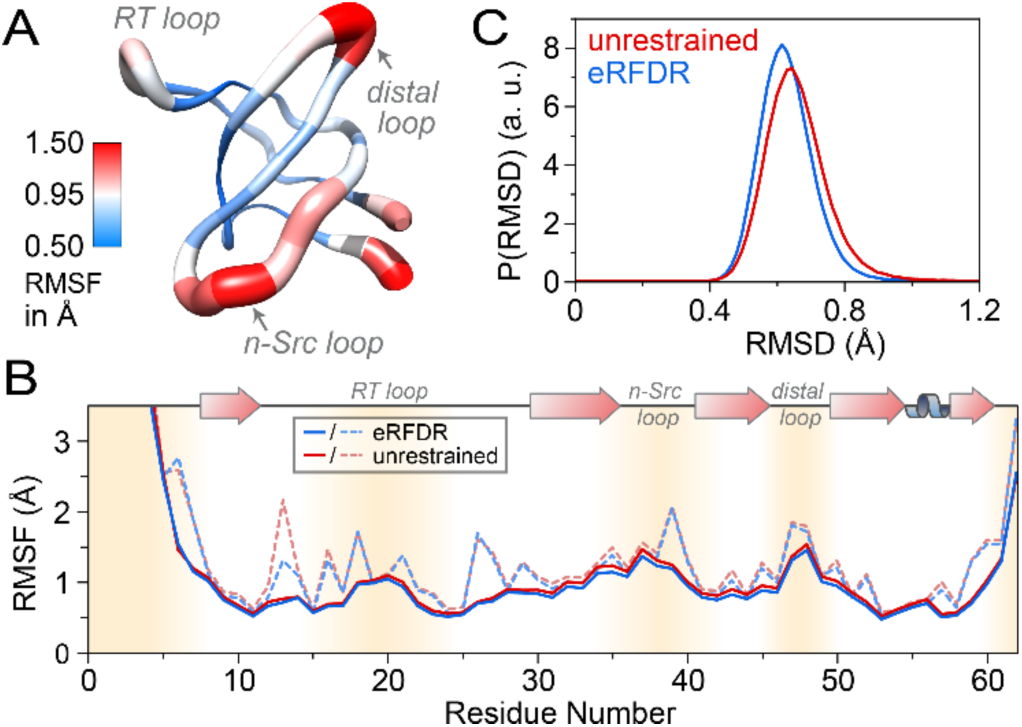
Restrained-ensemble MD simulations based on experimental eRFDR restraints. **A)** Structure of SH3 with C^α^-RMSF indicated as ribbon thickness. **B)** Comparison of the ensemble-averaged sequential C^*α*^ - and heavy-atom RMSF (solid and dashed lines, respectively) of eRFDR-restrained MD simulations (blue) to those from unrestrained MD simulations (red). **C)** Normalized probability distribution of C^α^-RMSDs for residues 7 – 61 with respect to the X-ray structure 2nuz after energy-minimization and equilibration, comparing unrestrained MD simulations (red) and an ensemble of MD simulations restrained by exact solid-state restraints (blue). All restraints accord to eRFDR data from the fully protonated sample.

Here we have shown that high-accuracy ssNMR distance restrains for solid proteins for integrated structural biology are afforded by combining fully protonated proteins, facilitated at MAS rates in the 100 kHz regime, and time- and context-dependent assessment of ^1^H-^1^H magnetization transfer. Adding to the low sample amount required (a single 0.5 mg non-deuterated sample), the procedure, which has also been made available via the Bruker “Dynamics Center” in a semi-automated fashion, enables very high numbers of accurate restraints, outcompeting the advantages of partially deuterated samples, and allows for ssNMR-restrained ensemble MD to elucidate experiment-driven spatial dynamics in solid protein samples at atomic detail. For the small globular protein used as a case example in this proof of concept study, the protein dynamics are confirmed to be highly similar between solution and crystal. Conversely, for manifold future targets, including supramolecular assemblies, fibrils, and membrane proteins, whose plasticity is functionally modulated by the complexation/aggregation state, ssNMR-restrained ensemble MD simulations will be key to elucidate diverse biological questions, possibly also addressing the challenge of in-silico reconstruction of the underlying lattices^[21]^.

## Acknowledgements

Financial support is acknowledged from the Deutsche Forschungsgemeinschaft (DFG, German Research Foundation) in the context of SFB 749, TP A13 (project number 27112786), SFB 1309, TP 03 (project number 325871075), and the Emmy Noether program. This work was funded under Germany’s Excellence Strategy – EXC 2033 390677874 – RESOLV and EXC 114 – 24286268. The authors gratefully acknowledge the computing time provided on the Linux HPC cluster at Technical University Dortmund (LiDO3), partially funded in the course of the Large-Scale Equipment Initiative by the German Research Foundation (DFG) as project 271512359.

Note: The pipeline has been integrated (called “Exact SolidState Distances”) in the Bruker Dynamics Center.

